# Differences in layer-specific activation of the insula during interoceptive vs. exteroceptive attention

**DOI:** 10.1101/2025.10.08.679044

**Authors:** Matthias Müller-Schrader, Frederike H. Petzschner, Jakob Heinzle, Lars Kasper, Katharina V. Wellstein, Johanna Bayer, Maria Engel, Samuel Bianchi, Lilian A. Weber, Klaas P. Pruessmann, Klaas Enno Stephan

## Abstract

Attention modulates the relative weighting of information and plays a critical role in modern theories of perception. While its role in exteroceptionthe perception of the external environmenthas been extensively studied, attentional processes in interoception, the perception of bodily states, are less well understood. In this study, we exploited high-field (7 Tesla) functional magnetic resonance imaging and concurrent field monitoring to investigate interoceptive attention at the level of cortical layers during a heartbeat attention task.

Voxel-wise analyses reveal increased activity in the bilateral dorsal mid-insula during interoceptive attention (attending to one’s heartbeat) compared to exteroceptive attention (attending to auditory white noise). Layer-specific analyses further demonstrate that this difference in activity is significantly higher in upper compared to lower cortical layers. This activation pattern persists after accounting for potential vascular artifacts through a deconvolution analysis with a physiological point spread function (PSF).

To our knowledge, these findings represent the first empirical demonstration of layer-specific processes during interoceptive attention in human cortex. They may prove useful to inform and constrain theories of computational principles and physiological implementation of interoception.

## Introduction

Perception is not only confined to the processing of external sensory inputs, such as sound, vision or proprioception (exteroception). It also encompasses the processing of signals originating from within the body – a domain known as interoception (Craig, 2002, 2009; Khalsa et al., 2018; Petzschner et al., 2022; Toussaint et al., 2024). The capacity to infer on the states of internal organs, such as heart and lungs, is indispensable for maintaining homeostasis deploying and allostatic controls, the physiological and adaptive processes that underpin survival.

Given the central role of interoception for monitoring and regulating the body’s internal milieu, it follows that disruption can have far-reaching consequences. Indeed, impairments in interoceptive processing have been linked to a broad spectrum of somatic and psychiatric disorders, including depression, anxiety, eating disorders, chronic fatigue, and chronic pain (Paulus and Stein, 2006; Paulus et al., 2016; Stephan et al., 2016; Seth and Friston, 2016; Petzschner et al., 2017; Khalsa et al., 2018).

Historically, interoception and exteroception have been treated as distinct domains, and studied separately. However, a wave of recent theoretical advancements has emphasized the need for a more integrated, mechanistic understanding of brain-body interactions. At the heart of this shift lies the proposition that interoception and exteroception may operate on shared algorithmic principles. Specifically, hierarchical Bayesian (or predictive processing) theories such as predictive coding (Rao and Ballard, 1999; Friston, 2005), initially developed to explain exteroceptive perception, have recently been extended to the interoceptive realm (Seth et al., 2012; Seth, 2013; Barrett and Simmons, 2015; Seth and

Friston, 2016; Petzschner et al., 2017, 2021). These theories posit that the brain actively constructs a generative model of sensory inputs – whether arising from the external or internal (bodily) world – and employs this model to infer the body’s current state within its environment. This model is continuously refined through prediction errors, i.e. mismatches between anticipated sensations and incoming sensory signals. The influence of these prediction errors, in turn, is weighted by their precision, a measure of relative uncertainty (precision-weighted prediction errors). Importantly, these computational theories postulate which computational quantities (predictions, prediction errors and precisions) are required for interoception, but, as explained below, have also been extended to where these quantities are computed in the brain, including the role of cortical layers.

Attention occupies a central position in predictive processing frameworks, serving as a key mechanism for modulating the influence of sensory information on perceptual inference (Friston, 2009; Hohwy, 2012). Specifically, directing attention to a particular sensory channels is thought to enhance the precision of its signals, thereby amplifying the impact of the prediction errors it conveys (Feldman and Friston, 2010). Disruptions in this process – manifested as imbalances in precision weighting or salience assignment – have been proposed as fundamental mechanisms underlying psychiatric disorders within predictive coding accounts (Adams et al., 2013; Friston et al., 2014; Lawson et al., 2014; Quattrocki and Friston, 2014; Haker et al., 2016; Stephan et al., 2016).

Despite its clinical significance, assessing attentional effects presents significant challenges, particularly in the interoceptive domain. Unlike exteroceptive signals, many internal bodily signals are not directly accessible to conscious awareness and are inherently difficult to manipulate in controlled experimental settings. In particular, measuring attentional effects of interoception requires readouts of neural activity that can capture subtle changes in processing of sensory inputs from the body. In previous work, we used EEG to explore the interaction between attention and the heartbeat-evoked potential (HEP) (Petzschner et al., 2019). In that study, we found a significant modulation of the HEP during interoceptive attention compared to exteroceptive attention, particularly within the 524 ms to 620 ms time window after the R-peak (Petzschner et al., 2019). Moreover, this attentional effect correlated with self-reported autonomic reactivity. The results aligned with prior research suggesting that attention can influence heartbeat-evoked potentials (Schandry et al., 1986; Montoya et al., 1993; García-Cordero et al., 2017).

Predictive coding frameworks of interoception have proposed that different regions and cortical layers of the insula play important roles (Barrett and Simmons, 2015). The insula, can be roughly divided into three parts that are distinguished cytoarchitectonically, particularly with the regard to the degree of how strongly the granular layers (layer IV) is expressed. The ventral portion of the anterior insula is agranular, while the dorsal extension towards the posterior insula is granular and the middle section is dysgranular (Mesulam and Mufson, 1985; Morel et al., 2013). More generally, anatomically extended versions of hierarchical Bayesian theories of perception (Friston, 2008) hypothesize that prediction error signals are primarily computed in the supragranular layers (I-III) Additional neurons in the neighbourhood (in particular in layers II and III in the same cortical column) are thought to function as precision units, which change the gain of the prediction-error computing neurons dynamically. These signals are then projected to deeper layers, particularly layer V, where predictions are thought to originate (Friston, 2008; Shipp et al., 2013; Barrett and Simmons, 2015).

While the relatively low spatial resolution of EEG makes it challenging to pinpoint the precise locations of cortical activity, recent advances in functional neuroimaging, particularly lamina-specific fMRI investigations (“layer fMRI”), offer a promising approach for investigating the anatomical hypotheses related to the insula’s contribution to interoception (Stephan et al., 2019). Layer fMRI exploits submillimeter resolution to to study brain activity non-invasively at the level of cortical layers (Polimeni et al., 2010; Goense et al., 2012; Muckli et al., 2015; De Martino et al., 2015; Kok et al., 2016; Petridou and Siero, 2019; Lawrence et al., 2019).

It has been successfully applied in the exteroceptive and motor domain, including studies of attentional modulation in visual and auditory processing: (De Martino et al., 2015; Klein et al., 2018; Gau et al., 2020; Liu et al., 2021; van Mourik et al., 2023; Heynckes et al., 2023).

In contrast to several applications of laminar fMRI in the exteroceptive domain, to date we are not aware of a layer fMRI study that studied interoceptive attention (or other aspects of interoception), leaving a critical gap in the literature. Here, we use a heartbeat attention task in combination with layer fMRI to explore the impact of interoceptive attention on cortical-layer-specific activity within the insula. In this task, adapted from Petzschner et al. (2019), participants switch their attention between an exteroceptive stimulus (auditory white noise) and an interoceptive stimulus (their own heart-beat). Importantly, the task controls for stimulus-related confounds and attentional demands per se, and enables one to isolate interoceptive vs. exteroceptive attentional effects (see Petzschner et al. (2019) for details).

Futhermore, we exploited high-field MRI (7 Tesla) in combination with concurrently field-monitored spiral readout trajectories and image reconstruction by the inversion of an advanced signal model (Barmet et al., 2008; Wilm et al., 2012; Engel et al., 2018; Kasper et al., 2022) in order to obtain functional MRI images with 0.9 mm in-plane resolution. The statistical analyses were pre-specified in a preregistered analysis plan (Müller-Schrader et al., 2023). By leveraging the high spatial resolution of layer fMRI, this study aimed to provide novel insights into the neural mechanisms underlying attentional modulation in the interoceptive domain, advancing our understanding of predictive processing in bodily perception.

## Results

In order to assess lamina-specific attentional processes during interoceptive attention, we analyzed high-resolution fMRI data (0.9 mm in-plane resolution) from 19 participants performing a heartbeat attention task (Petzschner et al., 2019). In this task, participants alternated between attending to their own heartbeat (interoceptive attention) and a continuous white noise stimulus (exteroceptive attention) in 16-second blocks. The task was carefully designed to isolate the effects of attentional focus on neural activity while avoiding confounds from differences in sensory input of task structure (Petzschner et al., 2019) (see Methods).

### Non-layer analysis

First, we performed a classical analysis, which does not incorporate layer-specific information, to asses the effect of interoceptive attention on activity in our a priori region-of-interest (ROI), the insula (Figure 2). Insular cortex was localized using an anatomically defined mask based on the Brainnetome atlas (Fan et al., 2016).

An *F* -test of the contrast *c*_1_ = *attention to heart attention to sound* revealed one significant cluster in the insular mask of each hemisphere. Using cluster-level inference (*p* < 0.05 FWE with cluster defining threshold *p* = 0.001), we identified a cluster in the left insula (left: *F* = 64.4 with peak at (*−*36, 2, 6); MNI space) and another in the right insula (*p*_*peak,F W E*_ *<* 0.001*/F* =81.4 with peak at (46, 4, 4)). Both clusters predominantly encompass the dorsal mid-insula in their respective hemispheres, likely corresponding to a part of the dysgranular insula (see Figure 2 and Supplementary Figure S1).

### Layered attention effect

We next examined the effect of interoceptive attention across different layers of the insula subregions. Of the 2 *×*6 insular subregions defined by the Brainnetome atlas, three subregions (L_dIg, L_dId and R_dId) had sufficient coverage by (out-of-sample) functional masks, defined as containing at least 10 voxels per subject and layer in that region (masks are different for each subject). Consequently, the analysis of layer-specific attention effects, using ANOVA to test for attention (heart vs. sound) x layer, was conducted within these three regions only. The interaction term was significant in L_dId and R_dId (L_dIg: *p* = 0.171; *F*_1,18_ = 2.03, L_dId: *p* = 0.001; *F*_1,18_ = 15.63 and R_dId *p* = 0.03; *F*_1,18_ = 5.55).

However, after correcting for multiple comparisons, only the interaction term for the left dysgranular insula (L_-dId) remained significant. For the significant interaction in L_dId, effect sizes for contrast *c*_1_ were *d* = 1.0 in lower layers and *d* = 1.45 in upper layers. See Figure 3 for a visualisation. Full ANOVA tables and additional details are provided in the Supplementary Material (Table S1 - S3).

### Layer profiles

Figure 4 illustrates the layer profiles (of parameter estimates from the general linear model) for interoceptive attention to the heartbeat (red) and exteroceptive attention to the sound (blue). Notably, the profiles show values around 0 at the white matter/gray matter (WM/GM) boundary. While there is asymmetry between hemispheres, the profiles exhibit more pronounced variations across insular subregions than between hemispheres. Except for the dorsal agranular insula (*dIa*), profiles have either a monotonic increase towards the GM/CSF boundary or remain around 0. Single-subject profiles can be found in Supplementary Figure S4.

Strikingly, in the dorsal granular insula (*dId*), there is a clear separation between the profiles for attention to heartbeat and sound, with a possible distinction also observed in the dorsal agranular insula (*dIa*). For the remaining insular subregions, the profiles for both conditions are similar.

### Deconvolution of layer profiles

A general challenge for interpreting layer fMRI results is that BOLD signal in upper versus lower layers can be increased by venous blood draining effects (Uludağet al., 2009; Koopmans et al., 2011; Heinzle et al., 2016; Markuerkiaga et al., 2016). This potential confound might explain the monotonic increase of signal towards the pial surface observed in Figure 4. In order to address this issue, we applied a previously proposed deconvolution strategy (Markuerkiaga et al., 2021a); this analysis was post-hoc (i.e. not included in our ex ante analysis plan). Specifically, we resampled the contrast values onto layers of the same cortical depth using a spatial GLM and deconvolved the results using a physiological point-spread function, which accounts for potential draining effects. The resulting profiles are displayed in Figure 5. The first column on each side shows the raw (binned) profiles, the second column presents the resampled profiles from the spatial GLM and the third column displays the deconvolved profiles.

An repeated-measures ANOVA, with factors for attention condition and layers, revealed a significant interaction between attention and layer in the left dysgranular insula (L_did, *p* = 0.003) and the right dysgranular insula (R _did, *p* = 0.029; both uncorrected).

This suggests that the observed interactions between attention condition and layer are not simply explained by draining effects.

## Discussion

To our knowledge, this study represents the first application of layer fMRI to interoceptive processing and the first use of non-Cartesian readout trajectories with concurrent field monitoring for layer fMRI.

Using a carefully controlled heartbeat attention task (Petzschner et al., 2019) and 7 Tesla MRI with with non-Cartesian readout trajectories and concurrent field monitoring for layer fMRI, we obtained high-resolution layer fMRI data (0.9 mm resolution) of attention to heartbeat versus attention to auditory stimuli. We found that attending to one’s heartbeat elicited stronger activation than attending to an external auditory stimulus, with significant activation clusters in the dorsal mid-insula of both hemispheres (“dorsal dysgranular insula” or area dId, accourding to the Brainnetome atlas). A layer-specific analysis further revealed that this attentional effect was stronger in upper cortical layers than in lower layers, suggesting a potential laminar differentiation in interoceptive processing.

### Interoceptive Attention in the dorsal dysgranular insula

Our findings align with previous fMRI studies (without analysis of layer-specific activations) reporting interoceptive attention effects in the mid and posterior insula. Notably, Simmons et al. (2013) identified two activation sites in the right mid-insula using an interoceptive attention paradigm that combined attention to both the heart and the stomach. One of these sites, located in the dorsal mid-insula (peak coordinates (37, *−* 3, 16)_*T*_ and (45, 1, 0)_*T*_), closely coincides with our observed peak activation in the right dorsal dysgranular insula. When comparing coordinates, this peak lies within approximately 5 mm of our observed peak activation, suggesting a strong spatial overlap. In contrast, the other activation peaks reported in Simmons et al. (2013) are more than 12 mm away (see Supplementary Material and Figure S6 therein for details).

Similarly, Avery et al. (2015) reported activation in the bilateral mid- insula regions (peak coordinates (− 30, 1, 16)_*T*_ and (37, 1, 17)_*T*_), as well as in the right dorsal posterior insula (34, *−*14, 20)_*T*_) during an interoceptive attention task (interoceptive condition: heart, stomach and bladder). Avery et al. (2014) further identified group-level differences in the bilateral dorsal mid-insula, with peak coordinates at (39, *−*6, 16)_*T*_ and (*−*34, *−*6, 16)_*T*_. These activations in the mid-insula are qualitatively consistent with our observations.

Beyond studies focusing on cardiac interoception, fMRI research has also examined interoceptive attention to respiration. Farb et al. (2013) reported bilateral activation in the posterior insula (peak coordinates: (33, *−*18, 21)_*M*_ and (*−*33, *−*15, 21)_*M*_), as well as in the right mid-insula (33, *−*6, 18)_*M*_ during breath-focused interoceptive tasks. These activation sites are positioned more dorsally than the clusters observed in our study, suggesting that interoceptive attention to different physiological signals, such as heartbeat versus breathing, may recruit distinct but overlapping regions of the insula. This is consistent with animal studies showing that different interoceptive signals project to spatially separated subregions of the insula (Evrard, 2019). However, these differences may also reflect variations in cognitive or attentional demands when monitoring respiratory versus cardiac signals.

Wang et al. (2019) further reported activation in what they termed bilateral anterior insula (peak coordinates: (34, 20, 4)_*M*_ and (*−*30, 20, 8)_*M*_). As shown in Supplementary Figure S6, the first activation peak partly overleaps with one of the activation clusters observed in our study, whereas the second peak is located a few millimeters from the corresponding cluster in the opposite hemisphere.

In a recent study, Adamic et al. (2024) report a bilateral overlap of activation in the dysgranular mid-insula for an interoceptive attention task (interoceptive condition: attention to heartbeat, breathing and stomach) and link hemispheric differences in that overlap to anxiety, depression and eating disorders.

This suggests that while both heartbeat, stomach and breath-focused interoception engages common regions of the insula, subtle spatial distinctions may reflect **modality-specific computations**. Future studies that directly compare heartbeat and breathing-related interoception within the same individuals using high-resolution fMRI could further clarify the functional organisation of the insula in interoceptive processing.

### Layered analysis of interoceptive attention

The use of spiral readout trajectories allowed us to achieve a nominal fMRI resolution of 0.9 mm, a level of precision commonly considered sufficient for layer-specific fMRI investigations (Finn et al., 2021). Here, for the first time, we applied non-Cartesian readout trajectories with concurrent field monitoring for layer fMRI. This approach requires offline reconstruction of MR images using the inversion of an advanced signal model, which accounts for coil sensitivities and static and dynamic off-resonances (Kasper et al., 2022). Importantly, this method offers improved geometric accuracy, making it particularly advantageous for layer fMRI applications.

To examine the layer-dependent effects of interoceptive attention, we used functional masks of the lower layers (25 % relative cortical depth) and upper layers (75 % relative cortical depth) within anatomically defined insular subregions. A repeated-measures ANOVA revealed a significant interaction between layer (upper/lower) and attention condition (heartbeat/sound) in the left dorsal mid-insula (area dId, according to the Brainnetome atlas; *p* = 0.001, *F*_1,18_ = 15.63). The nature of this interaction means that the BOLD signal difference between upper vs. lower cortical layers is greater for interoceptive attention than for exteroceptive attention (Figure 3 and Supplementary Figure S4). The interaction observed in the equivalent area in the right hemisphere (*p* = 0.03*/F*_1,18_ = 5.55) did not survive multiple comparisons correction.

In order to address the important potential confound of blood draining effects affecting preferentially BOLD signal in the upper layers of cortex, we used a previously proposed deconvolution approach (Markuerkiaga et al., 2021a). This analysis suggested that the observed differences between upper and lower layers do not simply reflect a haemodynamic artefact but reflect real differences in neuronal activity. As illustrated in Figure 3, the interaction effect in the left dorsal mid-insula (area dId) shows that compared to the exteroceptive (auditory) condition, interoceptive attention elicits a stronger signal increase in upper cortical layers than in lower layers. This suggest that the significant activations found by our conventional voxel-wise (non-layered) analysis in this region are primarily driven by activation in response to heartbeat-focused attention rather than suppression during exteroceptive attention.

### Interpretation in the context of predictive coding theories

Our findings can be interpreted within the framework of predictive coding theories of interoception (Barrett and Simmons (2015), Petzschner et al. (2021), but see discussion on limitations). According to these models, interoception relies on hierarchical Bayesian inference, where the brain continuously generates top-down predictions about internal bodily states and integrates them with bottom-up sensory inputs from the body.

In this framework, upper cortical layers (supragranular layers) are thought to play a dominant role in processing prediction errors, while deeper layers (infragranular layers) are more involved in generating top-down predictions (Friston, 2005, 2008; Shipp et al., 2013). Our findings of greater interoceptive attention effects in upper layers of the dorsal mid-insula (area dId) may therefore reflect an enhanced weighting of ascending sensory prediction errors when focusing on internal bodily signals.

One possible interpretations is that interoceptive attention modulates the precision of ascending sensory information, increasing the gain of prediction errors from visceral afferents. This is consistent with prior propos als that attentions serves as a mechanism for precision weighting in predictive coding, selectively amplifying the influence of prediction errors from attended stimuli (Feldman and Friston, 2010). Our findings align with this view by suggesting that interoceptive attention does not simply increase overall insular activation but selectively alters the balance of activity across cortical layers.

### Technical considerations and potential limitations

Several technical considerations must be taken into account when interpreting our findings.

While our results demonstrate that interoceptive attention increases the BOLD signal more strongly in the upper layers of the dorsal mid-insula (area dId) compared to lower layers, this does not necessarily imply a stronger increase in neural activity in upper layers. This distinction is important due to the complex nature of hemodynamic coupling, which governs how neural activity translates into the measured BOLD signal. In particular, ascending and pial draining veins introduce a known bias toward stronger signals near the cortical surface, potentially distorting the true layer-specific activation patterns (Koopmans et al., 2011). Furthermore, gradient-echo acquisitions as used in the present work, are sensitive to macrovascular effects, particularly draining veins (Uludağet al., 2009). The observed effects could thus either reflect true layer differences in neural activity, with interoceptive attention preferentially increasing activity in supragranular layers or be influenced by blood draining effects.

The cortical depth profiles (Figure 4) offer a more detailed perspective to assess the potential impact of draining effects: The profiles for the dorsal mid-insula (are dId) clearly illustrate the key patterns identified in our analysis: a distinct separation between attention to heartbeat and attention to sound, consistent with the significant clusters in the *F*-map. Moreover, the profiles align with the ANOVA results, showing a near-zero effect (GLM parameter estimate) for attention to sound and a stronger increase in BOLD signal in upper than in lower layers for attention to heartbeat. However, a potential limitation arises from the monotonic increase in contrast values toward the gray matter (GM)/cerebrospinal fluid (CSF) boundary, which could potentially be attributed to blood draining effects. of layer-specific neural activity.

Having said, the profiles for other insular subregions (e.g. the “dorsal agranular insula”, according to the Brainnetome atlas) show a non-monotonic pattern that is not easily explained by simple vascular effects.

Several approaches have been proposed for disambiguating neuronal and vascular effects when interpreting layer fMRI results. One strategy is the spatial de-convolution approach introduced by Markuerkiaga et al. (2021b), which aims to correct for interlaminar leakage by modeling the measured activation profiles as a weighted sum of true laminar activity convolved with a layer-specific physiological point spread function (PSF). Since blood draining effects propagate only toward the pial surface, the convolution matrix has lower triangular form, making deconvolution more robust. Simulations from Markuerkiaga et al. (2016) further suggest that the PSF can be approximated with only two parameters: a peak value at a given layer and a tail value representing signal into upper layers, allowing for a simple experimental determination of correction factors.

An alternative approach involves dynamic causal modeling (DCM) of laminar activity (Heinzle et al., 2016). This method attempts to disentangle neural and vascular contributions by incorporating a hemodynamic delay parameter that accounts for differences in blood flow dynamics across cortical layers. However, this approach is better suited for event-related designs than for block designs (as used in this study), where sustained activa tion patters make it difficult to separate delayed vascular responses.

Finally, one could consider pial vein masking or phase regression techniques, which aim to reduce macrovascular BOLD contributions by leveraging phase data (Menon, 2002; Stanley et al., 2021).

In order to clarify then potential impact of draining effects, we conducted a deconvolution analysis using a physiological PSF (Figure 5). The results were qualitatively similar to our original findings. A repeated-measures ANOVA continues to show significant interaction between attention condition and layer in the left and right dorsal mid-insula (area dId). These results indicate that our observed intereaction effects are unlikely to be strongly influenced by draining artifacts, supporting the conclusion that interoceptive attention increases neural activity more strongly in upper layers than in lower layers.

In summary, we identified an effect of interoceptive cardiac attention in the bilateral dorsal mid-insular, with layer fMRI revealing significant laminar differences in the left dorsal mid-insula. Venous blood draining effects, which could potentially mimic layer-specific activation patterns, are unlikely to have significantly influenced this result, as indicated by a spatial deconvolution analysis. To our knowledge, these findings represent the first empirical demonstration of layer-specific processes during interoceptive attention in human cortex. They may prove useful to inform and constrain theories of computational principles and physiological implementation of interoception.

## Methods

The statistical analyses (hypothesis tests) of this study were specified *ex ante* in an analysis plan. (As explained in the Results section, one exception is the spatial deconvolution analysis which we decided to conduct after having obtained results on layer differences.) The analysis plan was written after image acquisition and reconstruction had been completed, but before conducting any form of data analysis. It was preregistered on Zenodo (Müller-Schrader et al., 2023).

The final analysis plan was based on two previous analysis plans. One concerns the layer analysis of an auditory mismatch study which was acquired with the same participants, but a different task (auditory mismatch paradigm). The other https://gitlab.ethz.ch/tnu/analysisplans/muellerschraderetal_hbatt_ohbm describes a preliminary non-layer group-level analysis (*F* -test) on the attention-to-heart dataset which was conducted on a subset (*N* = 16) of the participants, which was presented at OHBM 2023. That analysis plan was specified before performing any statistical analyses, afterwards, the non-layer group-level analysis was performed on all subjects reconstructed with sufficient quality so far. Note that the result of that *F* -test did not impact any of the choices for the subsequent analyses.

In the following, we will mostly follow the structure of the analysis plan (Müller-Schrader et al., 2023). If needed, we will make things more precise and also explain deviations from it.

### Participants

40 participants (age between 18 and 40 years) participated in this study, which was approved by the ethics committee of the Kanton of Zürich (BASEC-Nr.: 2016-01297). All participants gave their written informed consent before entering the study. Exclusion criteria included current medication (except hormonal contraceptives), past or present drug abuse, serious past or present brain disease, injury or surgery and aspects related to MR safety (having MRI-incompatible metal parts or fragments in the body, dependence on a hearing aid, claustrophobia and pregnancy) and MR quality (inability to lie still for a longer period).

Out of 40 participants, 19 were included in our final analysis. We obtained complete MR data from 29 participants, but had to exclude 10 (4 due to excessive movement/displacement, 4 because of shim problems precluding determination of appropriate *B*_0_ maps and 2 because of other problems with reconstruction) according to the exclusion criteria specified in the analysis plan. In two cases, participants reported in the debriefing after the experiment that they had heard the auditory white noise only on the right ear. These subjects were included in the analysis. In one case, the timing of the blocks had to be reconstructed from the task code, because the behavioral file including this information was missing.

The whole study, which also included other tasks, took place in two sessions. Between the sessions, participants filled in a set of questionnaires at home (not analysed here).

### Heartbeat Attention Task

Participants performed a heartbeat attention task (Petzschner et al., 2019) while lying in the MR scanner, white noise was played continuously on earphones.^1^ Instructions were shown using MR compatible video goggles (VisuaStim, Resonance Technology Inc., Northridge, CA).

The heartbeat attention task consisted of alternating 16- second blocks during which participants were instructed to direct their attention either to their own heartbeat (interoceptive attention, condition: *attention to heart*) or to a sound stimulus (exteroceptive attention, condition: *attention to sound*). Each block was followed by a rating period (maximum duration: 9 seconds (2 + 7 seconds)) and an inter-trial interval (ITI) with a variable length (714 seconds), so that rating and ITI took 16 seconds (see Figure 1).

**Figure 1.**
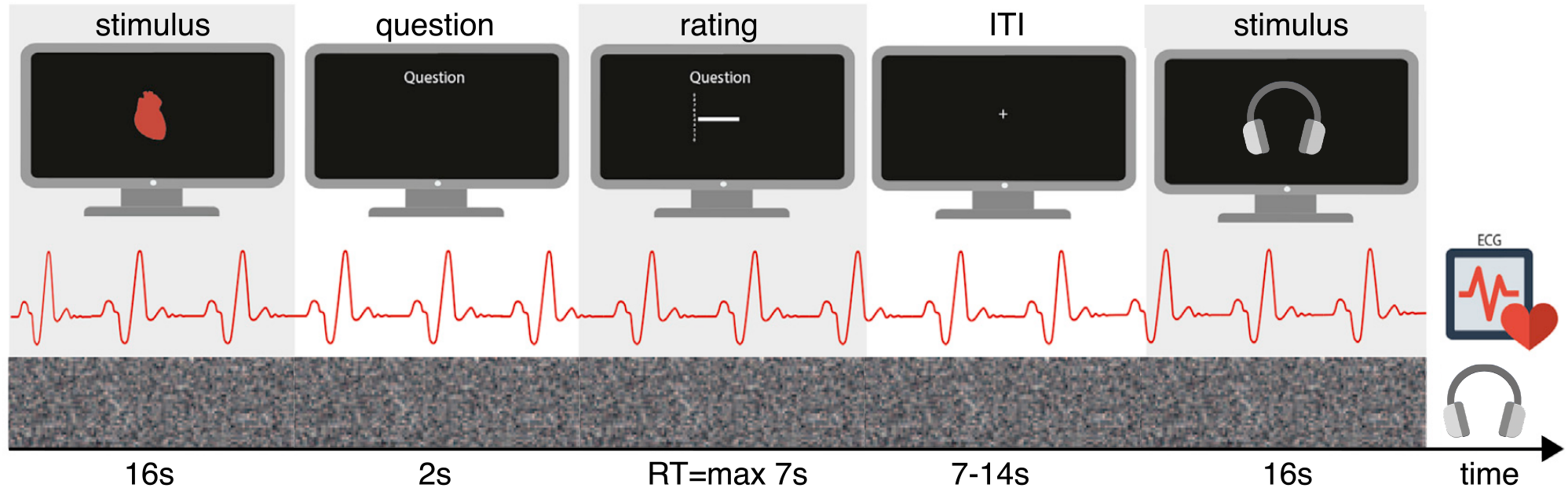
Structure of the heartbeat attention task. The task comprised 4 runs, each with 16 blocks, resulting in 32 blocks per condition and 64 blocks in total. Participants alternated between 16-second blocks of interoceptive attention (focusing on their own heartbeat) and exteroceptive attention (focusing on a continuous auditory stimulus, i.e. white noise). Following each block, a rating period (maximum duration: 7 seconds) required participants to answer a question about their experiences during the preceding attention block. The rating questions were included solely to maintain participants engagement throughout the task. This was followed by an inter-trial (ITI) of variable length. Adapted from Petzschner et al. (2019) under a CC BY-NC-ND license.

**Figure 2.**
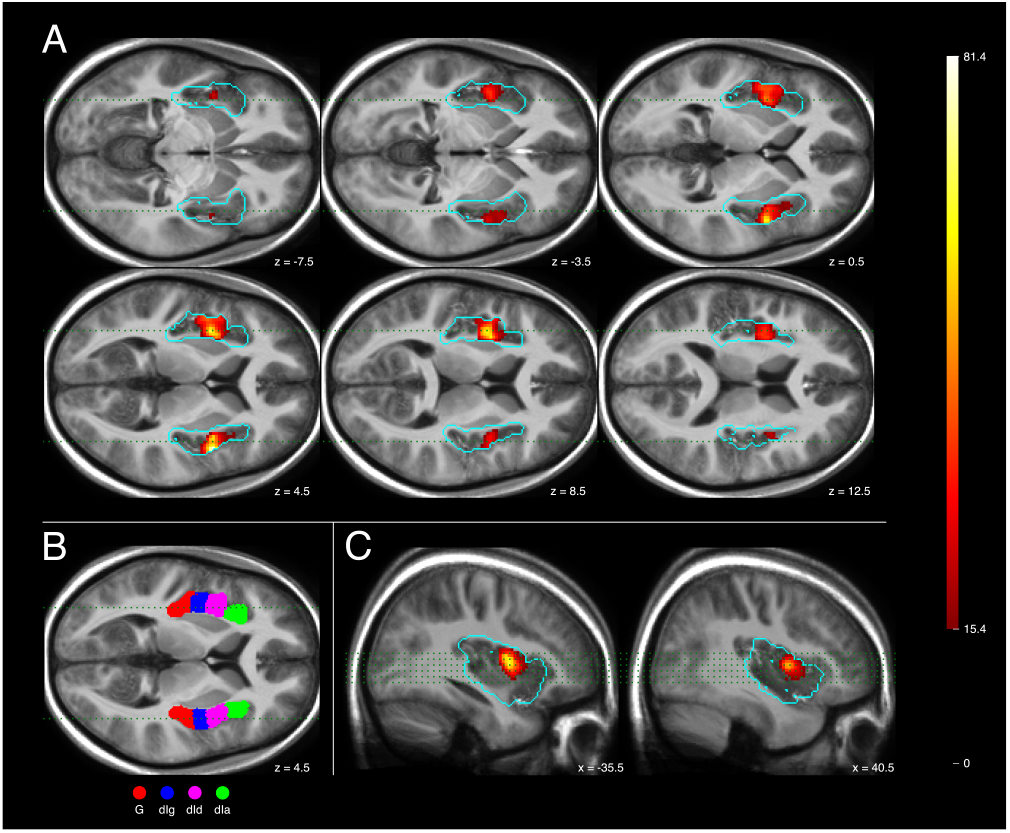
Classical Analysis of Interoceptive Attention Effects on Insula Activity. **A/C** (transverse/sagittal section*): Group level *F* -map of contrast *c*_1_ = *attention to heart − attention to sound*, thresholded at cluster level (*p* < 0.05, cluster defining threshold of *p* = 0.001). The cyan contour delineates the anatomical mask of the insula used for the analysis. The green dots indicate the locations of the sagittal (**A**) and /transverse section*s (**B**). **B** Insular mask with subregions: *G* hypergranular insula, *dIg*: dorsal granular insula, *dId* : dorsal dysgranular insula, *dIa*: dorsal agranular insula. Not visible on this slice are *vId_vIg*: ventral granular insula and *vIa*: ventral agranular insula.

**Figure 3.**
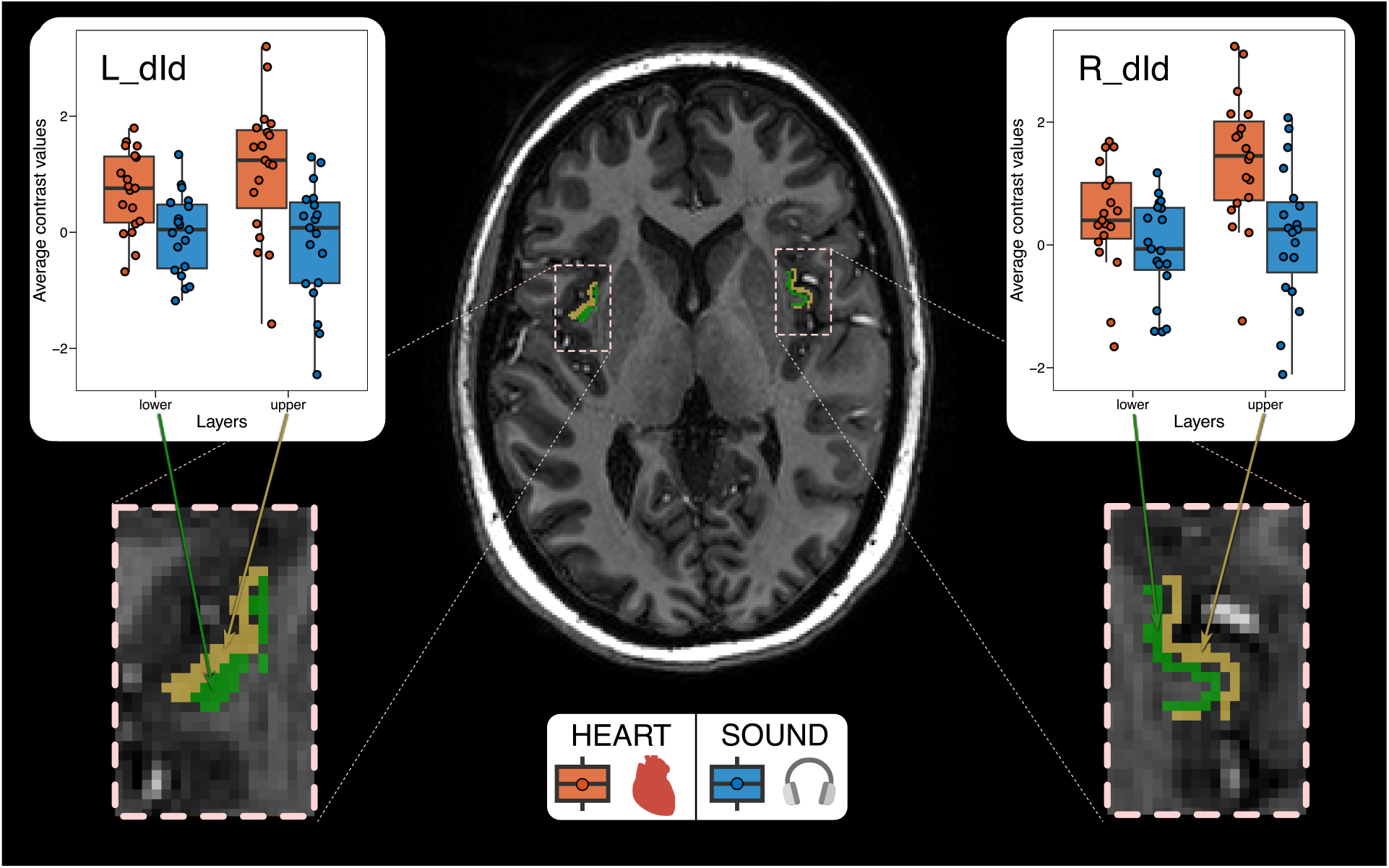
Visualization of within-subject ANOVA: Contrast values (red: attention to heart /blue: attention to sound) are averaged for each subject across the layer masks (green: lower layer at 25 % cortical depth /yellow: upper layer at 75 % cortical depth). On these values, which are shown in the plots for the left and right dorsal mid-insula (“dorsal dysgranular insula” or area dId, according to the Brainnetome atlas), a repeated measures (within-subject) ANOVA was performed. The interaction term *attention to heart/sound × layer* was tested for significance (L_dId: *p* = 0.005; *F*_1,18_ = 10.25, R_dId: *p* = 0.047; *F*_1,18_ = 4.54; L_dId remains significant after multiple comparison correction).

**Figure 4.**
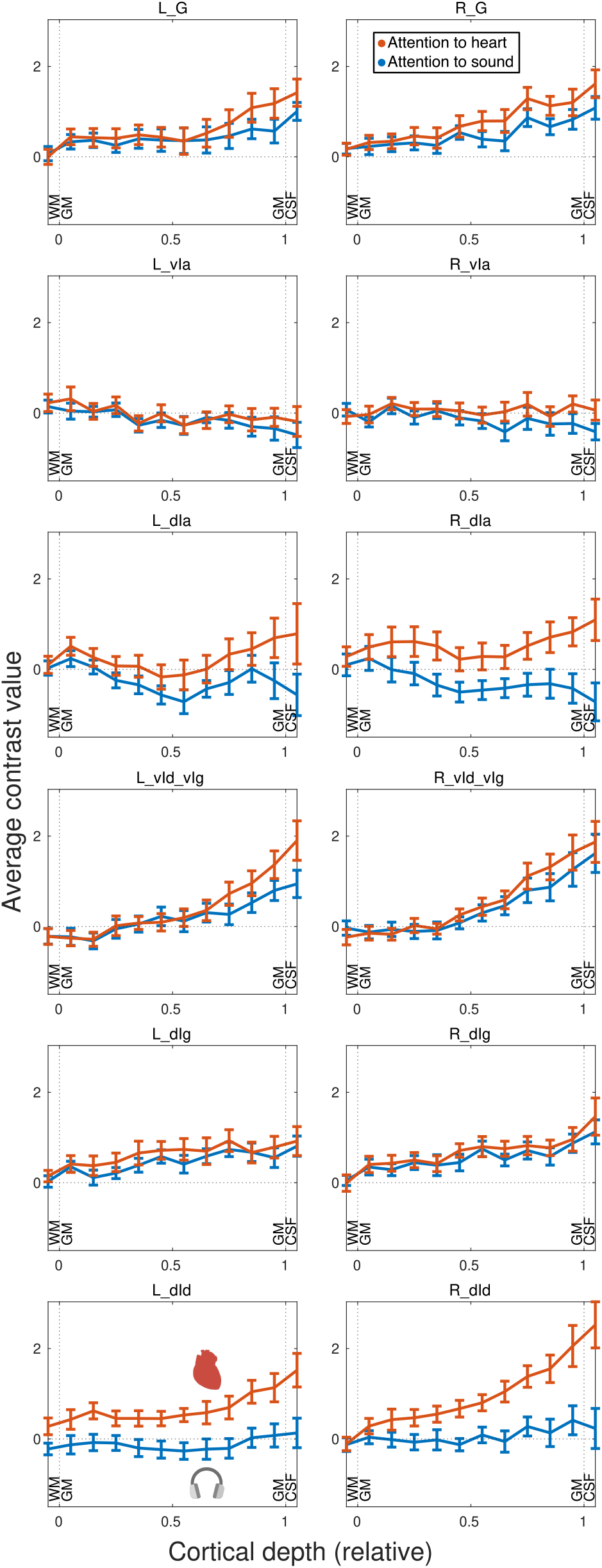
Layer profiles for parameter estimates from the GLM for interoceptive attention (*β*_attention to heart_, red) and exteroceptive attention (*β*_attention to sound_, blue). For each insular subregion, *βs* at the same cortical depth are averaged over subjects. The 15 % bin contains for example all voxels, whose center is located between the 10 % and the 20 %(equi-volume) relative cortical depth surface. Error bars indicate the standard error of the mean (over subjects). Please note that while the voxel size is 0.9 mm, the bin size for these profiles is around 0.2 mm to 0.4 mm, which introduced some smoothness in the signal. Region names (L/R indicate hemisphere): *G*: hypergranular insula, *vIa*: ventral agranular insula, *dIa*: dorsal agranular insula, *vId_vIg*: ventral granular insula, *dIg*: dorsal granular insula, and *dId* : dorsal dysgranular insula.

**Figure 5.**
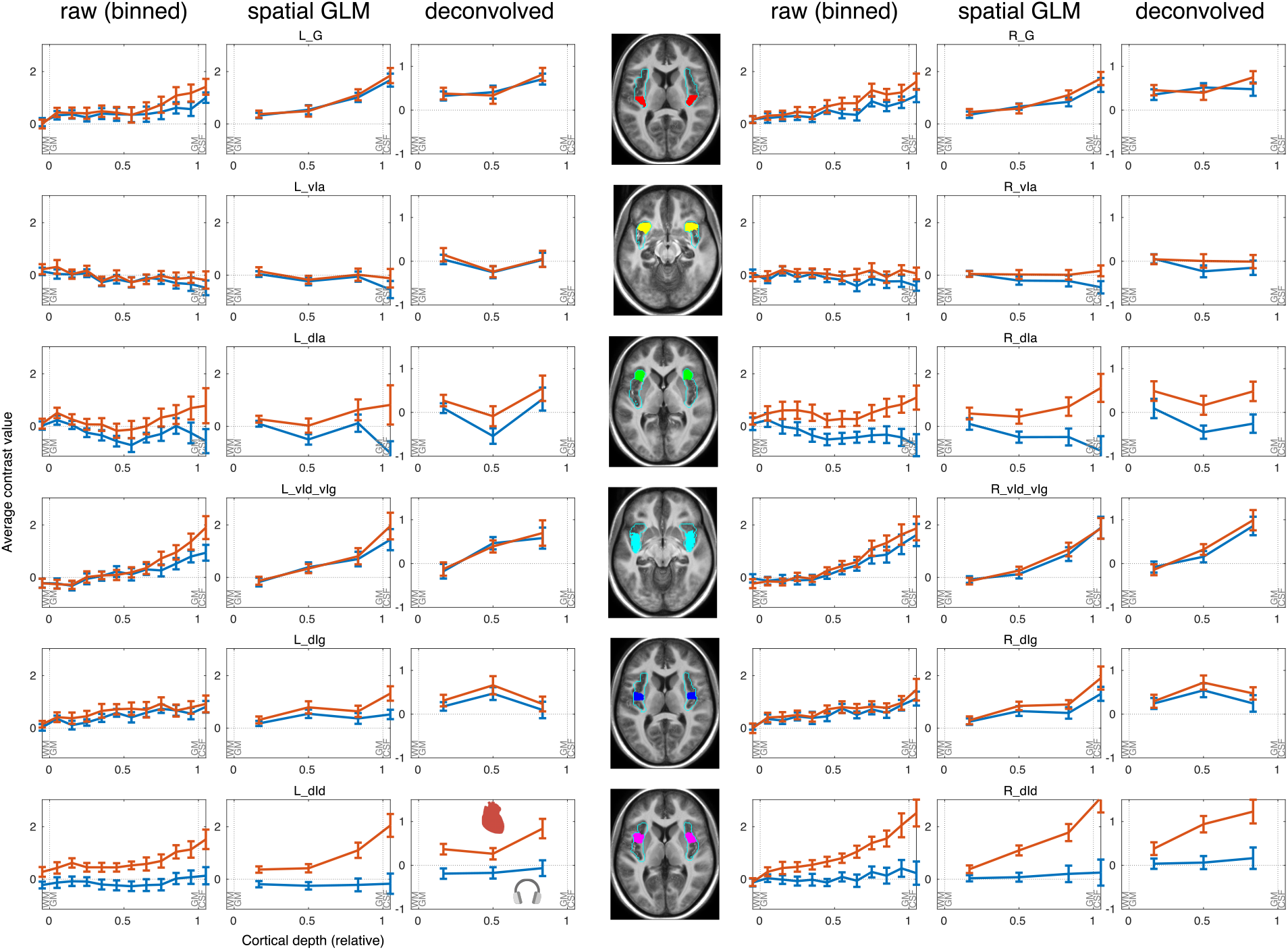
Results of deconvolution with a physiological PSF: The left column of each side shows the raw profiles for *β*_attention to heart_ (red) and *β*_attention to sound_ (blue), binned across cortical depth and averaged over subjects, as in Figure 4. The second column shows the profiles resampled by a spatial GLM onto three cortical and one CSF-layer (to account for pial veins). The third column shows the profiles for the cortical layers, which were deconvolved by a physiological PSF (*p*2*t*_3_ = 2.2, which corresponds to *p*2*t*_10_ = 6.3), that captures the draining effect. Each row represents one insular subregion (left and right hemisphere), the location of that area in MNI space is visualized in the middle (the *z*-coordinate is the center-of-mass). The error bars indicate the standard error of the mean (over subjects). A repeated-measures ANOVA showed a significant interaction *attention condition x layer* in the L_did (*p* = 0.003) and the R_did (*p* = 0.029, both uncorrected).

*Attention to heart* blocks were indicated by a visual heart symbol displayed on the screen for the entire block duration. Participants were instructed to focus their attention on their heartbeat and to notice any changes in heart sensations or heart rate, without physically measuring their pulse. *Attention to sounds* blocks were marked by a headphone symbol, which remained on the screen throughout the block. During these blocks, participants were instructed to attend to the continuous white noise played through headphones, noting any potential changes in the sound.

To ensure that any observed changes in brain activity reflected shifts in attention rather than differences in sensory stimulation, the white noise stimulus was played during the entire experiment. The volume was adjusted to ensure that it was clearly audible for all participants.

During both conditions, participants were prompted to rate specific aspects of their perception or associations with the previous block during the rating period. Questions varied across conditions (e.g., *How well were you able to concentrate on the white noise in the last block?* or *How much would you associate your perceived heartbeat in the previous block with the color red?*), with responses recorded on a scale from 1 to 10. These ratings served solely to maintain engagement and focus; they were not included in subsequent analyses.

The ITI was indicated by a fixation cross, during which participants were free to think about anything. Participants performed 4 runs of 16 blocks per run of the task, resulting in 32 blocks per condition in total.

### Data acquisition

The following description of our data acquisition procedure (next 4 paragraphs) is a near-verbatim copy of the description in our analysis plan (Müller-Schrader et al., 2023):

Participants were scanned on a Philips Achieva 7T MRI system (Philips Healthcare, Best, The Netherlands) using a quadrature transmit and 32-channel head coil (Nova, Wilmington, MA). In addition, 16 NMR-fieldprobes were mounted around the head (Dietrich et al., 2016), allowing for real-time monitoring of concurrent magnetic fields.

Functional images were acquired using a high-resolution single-shot spiral-out trajectory with 0.9 mm in-plane resolution (parallel acceleration *R* = 4, *T*_*E*_ =15 ms, 50 ms readout duration), combined in a multislice 2D sequence with 36 slices (0.9 mm slice thickness, 0.1 mm gap) and fat suppression module (SPIR), amounting to a volume TR of 3.128 s (similar to Kasper et al. (2022), but with a different readout duration/resolution). For each participant, we recorded four fMRI runs with 180 volumes each. Between runs 1 and 2, between runs 3 and 4 and after run 4, a fat-suppressed 2D Cartesian gradient-echo scan (*R* = 1) with six echos (echo times 6 ms, 7 ms,…, 11 ms) 1 mm in-plane resolution and a slice geometry equivalent to the functional images, was acquired (to compute *B*_0_-off-resonance and SENSE maps).

We acquired two anatomical reference images, one structural *T*_1_-weighted image with 0.8 mm isotropic resolution (Inversion recovery 3D Cartesian gradient-echo)^2^ and another 1 mm multi-echo Cartesian scan (*R* = 3) with whole brain coverage (120 slices). We will call these images *T*_1_ (image) and MEWB in the following. The *T*_1_ image has a contrast suited for Freesurfer’s surface reconstruction, but we chose the MEWB image as reference image because it was reconstructed the same way (including field monitoring) as the fMRI images. Finally, we acquired a 5 mm multi-echo scan (*R* = 1) with the same geometry as the MEWB image to calculate *B*_0_- and SENSE-maps for its reconstruction.

During scanning, heart rate was recorded using both, a pulse oximeter and ECG. Breathing was monitored using a breathing belt. Since the ECG data was too noisy for a reliable detection of peaks, we only used the data from the pulse-oximeter (and the breathing belt) for the correction of physiological noise.

### Reconstruction of MR images

Image reconstruction was done by inversion of an advanced signal model, which takes coil sensitivities and spatial and dynamic off-resonances into account (Kasper et al., 2022). Importantly, the correct reconstruction of fMRI images depends strongly on correct off-resonance maps. Inaccuracies in these maps – e.g. due to strong displacement of participants as result of movement – leads to blurring or signal loss in fMRI images. As specified in the analysis plan, we visually inspected the reconstructed fMRI images prior to any data analysis and excluded participants if we could not compute accurate off-resonance maps (shimming problems) or included nuisance regressors for volumes with blurring due to excessive displacement (more details below). We used an in-house MATLAB R2022a (The MathWorks, Natrick, MA, USA) implementation of that reconstruction algorithm, which also included an spike correction algorithm operating on the raw coil data.

### Preprocessing of data

Preprocessing of MR images was done using SPM12 (v. 7219) on MATLAB R2023a (The MathWorks, Natick, MA, USA).

We averaged the MEWB image over echos (RMS) and performed unified segmentation (Ashburner and Friston (2005)/spm_segment) on the result. The resulting bias-corrected image served as anatomical reference for the subject. All other images were coregistered to this image.^3^ Unified segmentation results in transformations (warping fields) that served to transform images between the subject spaces (defined by the individual MEWB-images) and the common (SPM-)MNI space. Note that all layer analyses were performed in the individual subjects’ space, avoiding distortions due to the nonlinear normalization.

To account for potential *T*_1_-saturation effects, we discarded the first 4 functional volumes of each fMRI run. The remaining functional images of each run were slicetiming corrected and realigned using SPM.

To determine the (affine) transformation between the functional space (defined by the first volume) and the anatomical subject space, we bias-corrected the mean functional image (again, using unified segmentation) and coregistered the result to the reference MEWB image. The resulting affine transformation was applied on (the headers of) all functional images.

The *T*_1_-weighted image was bias-corrected and coregistered to the reference MEWB image for each participant. Again, no reslicing was performed.

### Definition of anatomical masks

We used the Human Brainnetome Atlas (Fan et al., 2016) to define an anatomical mask of the insula in MNI-space (Region numbers 163 - 174). See the caption of Figure 2 for the region names. Using the deformation fields and the coregistration matrix, the masks were transformed to the functional space. To avoid cutting parts of the insula as consequence of imprecisions in normalization, we inflated the resulting masks by 2 voxels (=2 mm in MEWB space). The details of the procedure used to define subject specific masks are described in Appendix 1.

### Preparation of confound regressors

To account for confounds due to motion and physiological noise, we created the following nuisance regressors: Using the TAPAS PhysIO Toolbox R 2022b, V 8.2.0 (Kasper et al., 2017), which is part of the TAPAS software collection (Frässle et al., 2021), we created the following 16 physiological nuisance regressors based on pulse oximeter and breathing belt measurements:

- 14 RETROICOR regressors (Glover et al., 2000) of cardiac (3rd order) and respiratory (2nd order) phase and one interaction term (1st order)
- one respiratory volume per time regressor (RVT/Birn et al. (2008))
- one heart-rate regressor (to take into account effects due to heart-rate variability, see Chang et al. (2009))

To reduce the effect of motion-related artifacts, we included the 6 movement regressors obtained during realignment.

Furthermore, to take abrupt motion into account, we included a motion-censoring regressor for each volume with a frame-wise displacement (FD; this is the *L*_1_ norm of displacement, where angles are multiplied by 50 mm rad*−*1 as done in Power et al. (2012)) of 0.5 mm or more.

Finally, to compensate for blurring in the reconstruction due to the mismatch of the *B*_0_-map, we added additional nuisance regressors for each volume with a translational displacement of 3 mm or more compared to a reference volume.^4^ The reference volume was the first or last volume of the run dependent on whether the ME image to calculate the *B*_0_ map was acquired before or after the functional run, respectively.

Participants with one run (or more) with at least 36 nuisance regressors (≥20 % of the volumes of that run) were excluded from the analysis.

### Creation of layered surfaces

Grey matter surfaces (boundaries between grey matter/white matter respectively grey matter/CSF) were reconstructed with Freesurfer (v. 7.4.0-1 /Fischl (2012)). Following the recommendation in the Freesurfer wiki (Zaretskaya, 2024), we used *T*_1_-images in their original resolution, which were already bias-corrected in SPM (using unified segmentation, Ashburner and Friston (2005)) and entered them into Freesurfer’s automated preprocessing stream (using the recon-all command with the -cm option).

After ensuring the quality of surfaces by visual inspection, we smoothed the grey matter surfaces with Freesurfer’s mris_smooth command (with 1 iteration and 0 averages). Using surface tools (Wagstyl and Huth, 2023), we created surfaces at 25 % and 75 % cortical depth, which served as reference for infra-respectively supragranular layers. Cortical depth was thereby calculated following the equivolume-principle (Waehnert et al., 2014). In addition, we computed equidistance surfaces of cortical depth from 0 % to 100 % (in steps of 10 %) for the creation of cortical profiles.

The surfaces were then transformed to the subject fMRI space by applying the affine transformations obtained during coregistration. The same was done for the mask of the cortical ribbon (determined by Freesurfer during the reconstruction pipeline).

For each voxel within the logical conjunction of cortical ribbon mask and the insular mask (defined by the Brainnetome atlas), we calculated the signed distance to all surfaces within the same hemisphere (using the point2trimesh function /see Frisch (2015)). These distances formed the base for the following analyses.

### Extraction of layer masks

We defined the *lower layer* and *upper layer* mask to include all voxels, whose distances to the 25 % and 75 % cortical depth surface were ≤ 0.5 mm, respectively.

### Correction of insular layer masks

In order to compensate for possible coregistration or normalization errors, we inflated the anatomical insular masks by 2 voxels (≙2 mm). This reduced the risk of accidentally masking out parts of the insula but also led to the inclusion of other parts of cortex - especially the operculum opposite of the insula (compare Figure 6 B). While there is some variation in the exact microstructural border between operculum and insula, “the circular sulcus is […] an approximate border to the […] opercular area OP3 “ (Quabs et al. (2022), see also Eickhoff et al. (2006)). We hence shrunk the insular mask by keeping only voxels, whose lower layer was more medial than its upper layer. See Appendix 1 for details of the shrinking algorithm. This procedure successfully removed all voxels outside the circular sulcus (verified by visual inspection).

**Figure 6.**
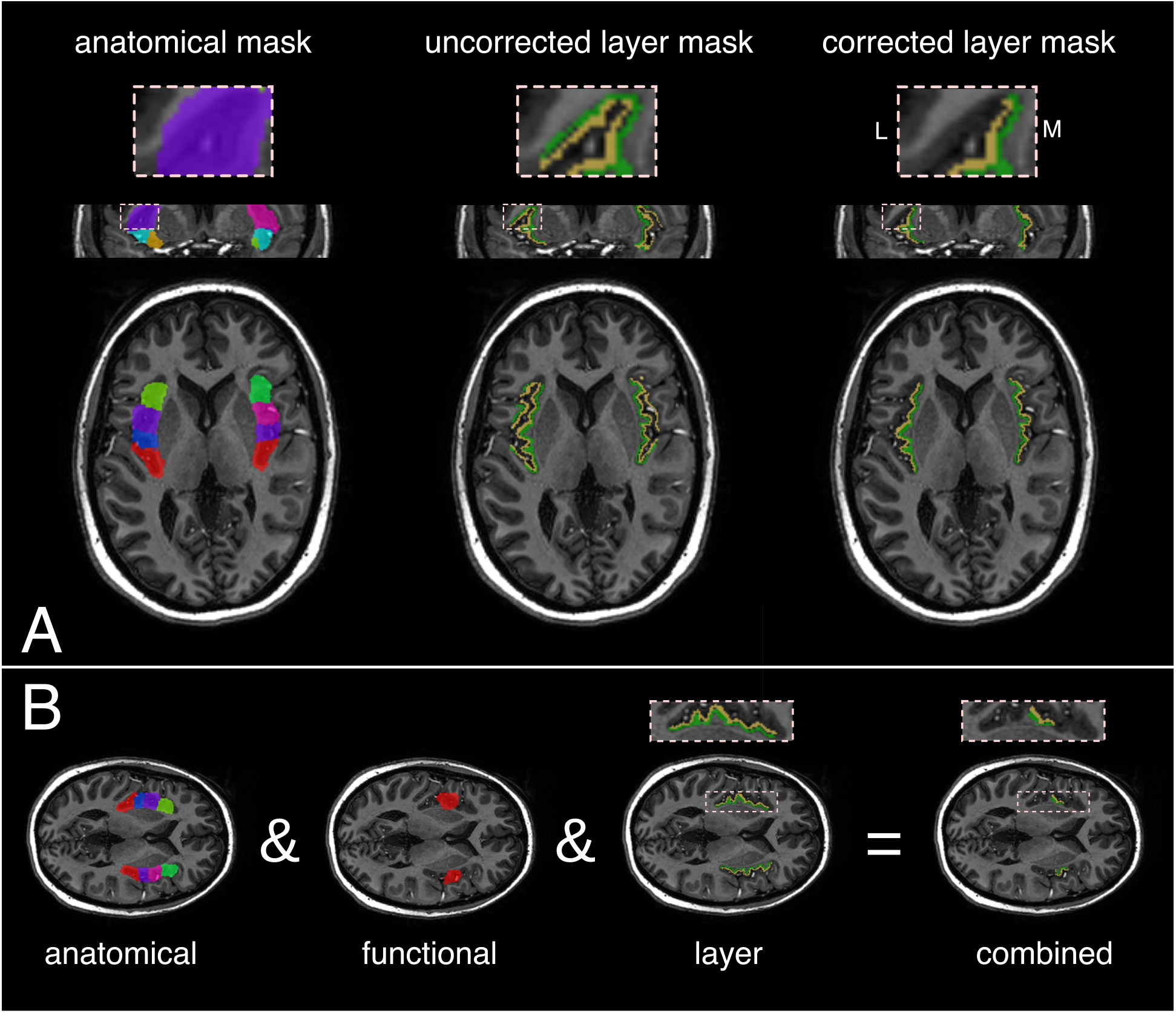
Details on mask computation. **A)** Correction of insular layer masks: The anatomical mask (**left**) was inflated by 2 voxels (=2 mm) to account for potential transformation imprecisions. The layer masks (**middle**) might hence also contain parts of the operculum - opposite of the insula. We correct for that (**right**) by only including voxels, whose lower layers (green) are more medial than their upper layer (yellow). *L* and *M* indicate *lateral* and *medial* orientation. **B)** Masking strategy for layer analysis: We combine an anatomical mask of the insula (with six subregions per hemisphere) with a functional mask (out-of-sample second level *F* -contrast) and a layer mask (25 %*/*75 % cortical depth).

### First-level GLM

For each subject, we constructed a first-level general-linear model (GLM) consisting of the block regressors *attention to heart* and *attention to sound* and the nuisance regressors described above (16 *physiological noise*, 6 *motion* and other *nuisance* regressors, as described above).

We were then interested in the contrast

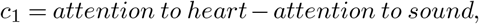

which represents the difference between attending to the interoceptive heart stimulus and to the exteroceptive sound stimulus.

### Classical second-level GLM and functional masking

For the classical(non-layer) group-level analysis, we transformed the contrast images (*c*_1_) into MNI space and smoothed them with a 6 mm FWHM Gaussian kernel.^5^

Within the insular mask, we then performed an (undirected) *F* -test to find regions that showed differential activation to the two attention conditions. Significance was tested at *p* < 0.05 family-wise error (FWE) corrected at the cluster level (cluster defining threshold *p* = 0.001) In addition, we computed effect sizes for significant activations in terms of Cohen’s *d*.

In order to increase sensitivity, we restricted the main layer analysis below to a subpart of the insula by a functional mask. While we had originally planned - as specified in the analysis plan (Müller-Schrader et al., 2023) - to use a mask obtained at the subject/first level (*F* -test on *c*_1_), a first analysis revealed that, for some subjects, a functional mask defined in this manner would contain no voxels. We therefore decided to deviate from the analysis plan and to compute the functional mask based on a second-level analysis. To avoid *double dipping*, we defined the mask for each participant out-of-sample, that is we computed the group activation for all but the participant in question. For example for the mask of subject 12, we combined data from subjects 1-11 and 13-19 at the second (group) level. For these subjects, we performed - similar to above - an *F* -test at the second level for the transformed contrasts *c*_1_ smoothed with a 6 mm FWHM Gaussian kernel.^6^ The results were FWE corrected at the cluster level (*p* < 0.05 with a cluster defining threshold of *p* < 0.001) to obtain the ROIs (mask), which were transformed back to the functional space of the individual (e.g., in the example above: participant 12).

### Creation of layer functional maps and layer second level analysis

We computed layered functional masks for each of the 2 *×* 6 subregions of the insula by combining (using logical *AND*) the anatomical mask of the insular subregion, the layer mask for infragranular/supragranular layers at 25 %*/*75 % cortical depth and the functional (out-of-sample group-level) mask, compare Figure 6.

Finally, to ensure that infragranular and supragranular masks contained the same parts of cortex, we reduced them in the following way: For each voxel in the *lower layer* mask, we computed the closest node in the corresponding lower surface (25 % cortical depth).^7^ For this node in the lower surface, we then considered the corresponding node on the upper surface (75 % cortical depth),^8^ and checked whether for any voxel in the *upper layer* mask its closest node in the upper surface was either the corresponding upper node or one of its neighbours. If this was not the case, we removed the voxel in the *lower layer* mask (which we started with). We repeated the same for all voxels in the *upper layer* mask.^9^ Within these layer functional masks, we calculated weighted averages over the regression weights *β*_attention to heart/sound_. Each voxel *v* was weighted by a weight *w*_*v*_ depending on its distance to the corresponding surface:

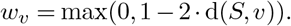

Here, d(*S, v*) denotes the distance between the voxel center and the corresponding surface (in mm). This weight was introduced so that voxels which were centered on the surface *S* received full weight, while voxels which were more distant received lower weight, until voxels, which were (more than) 0.5 mm away from the surface had weight 0.

We then analysed (for each insular subregion) the averaged *β*s (over space, within mask) in a within-subject-2-way ANOVA (also known as 2-way repeated measures ANOVA) with factors *upper layer* vs. *lower layer* and *attention to heart* vs. *attention to sound*. Our main research question was addressed by testing for a significant interaction using an *F* -test. To correct for multiple comparisons (3 tests in the 3 insular subregions covered by the functional masks)^10^, we applied Bonferroni-Holm correction to control the family-wise error rate below *p* < 0.05. For those regions, that showed a significant interaction, we computed layer-specific effect sizes

(Cohen’s *d*) for contrast *c*_1_. The ANOVA analysis was performed in R 4.4.1 (R Core Team, 2023) with rstatix (version 0.7.2, Kassambara (2023)). Plots were created with the tidyverse package (version 2.0.0, Wickham et al. (2019)).

### Layer profiles

To visualize the signal differences across cortical depth, we computed average values (over subjects and runs) in all cortical depths for *β*_attention to heart_ and *β*_attention to sound_, respectively, for each insular subregion. For this, all voxels inside the cortical ribbon mask were taken into account. Note that we reduced the mask as described in Appendix 1 to exclude potential contributions from operculum or other regions. This illustration of the layer activity was performed without any functional masking.

For each insular subregion and condition, we computed a histogram with bins centered at relative depths of

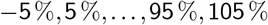

by using the surfaces at cortical depth^11^ of

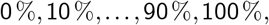

and averaging all *β*s, whose voxel centers lied between two successive surfaces (or outside of the 0 % or 100 % surface for the *−*5 % and 105 % bins). For example, for the 15 % bin, we averaged all *β*s, whose voxel center lies between the 10 % and the 20 % surface. These voxels were identified easily by the signed distances to the neighbouring surfaces.

### Profile-based classification of attention condition

In the analysis plan (Müller-Schrader et al., 2023), we had specified an additional out-of-sample analysis in case that “clear differences are observable in the layer profiles”. In that case, we would have trained a binary classifier on *z*-transformed layer single-subject profiles to predict the condition of the block (*attention to heart* /*attention to sound*) and tested it out-of-sample (training on all subjects except the test subject, averaging results across subjects). However, as seen in Figure S4 of the Supplement, the picture is not entirely clear. Therefore, we decided not to perform the out-of-sample classification analysis.

### Post-hoc analysis: Deconvolution of layer profiles

Since we did not observe any attentional effects in the dorsal mid-insula (area dId) during the attention-to-sound condition (compare Figure 3 and Figure 4), the observed interaction effect between attention condition and layer above could have different explanations: It could be a result of increased neuronal activity in upper layers or it could be explained by homogeneous neuronal activation throughout layers with apparent stronger BOLD activation in the upper layers due to blood draining effects.

We therefore decided go beyond our original analysis plan and conduct an additional post-hoc analysis, aiming to remove potential blood-draining effects based on a previously proposed deconvolution approach (Markuerkiaga et al., 2021a):

First, we performed a spatial unmixing of the data using a *spatial GLM* (Kok et al., 2016; Huber et al.,2017; van Mourik et al., 2019): Specifically, we defined the general linear model *Y* = *XB* + *ε*, where *Y* (dimensions *n*_Voxel_ *×* 1) denotes the data (i.e. single-subject regression coefficients *β*_*attention−to−heart*_ or *β*_*attention−to−sound*_, respectively), the design matrix *X* (dimensions *n*_Voxel_ *× n*_Layer_) encodes the distribution of layers across voxels and the noise *ε* is assumed to follow an (i.i.d.) normal distribution. Given this model, we obtained estimates *B* (dimensions *n*_Layer_ *×*1) of the contrast values in different cortical depths. We modeled 3 cortical layers (with identical thickness) to ensure that the thickness of these layers is greater or equal to the voxel size (note that the thickness of insular cortex is usually above 2.7 mm (Tanzer et al., 2021)). This reduction in the number of layers was chosen to avoid an overfitting of the spatial GLM as observed in Huber et al. (2017) and van Mourik et al. (2019). In addition, we included an additional layer to represent the possible contributions of pial veins in the CSF. The weights for the design matrix were then computed based on the partial volume of the layers within the voxel volume. To compute that partial volume, we used the normal vectors between voxel center and equal-cortical-depth-surfaces obtained earlier and computed analytically the partial volume of the cube cut by the plane defined by the normal vector. Note that this assumes that the curvature of the surfaces is small compared to the voxel size. Since van Mourik et al. (2019) showed that specifying a prior covariance structure on the noise results in “marginal improvements at best”, we used ordinary least squares (OLS) to invert the model.

On the spatially *unmixed* data *B* (restricted to the cortical layers), we then performed a deconvolution with a physiological point spread function. In Markuerkiaga et al. (2016) and Markuerkiaga et al. (2021b), it was shown in simulations and experimentally, that the draining through intracortical veins (perpendicular to the surface) can be described by a triangular convolution matrix, which has only one degree of freedom - the ratio *p*2*t* between diagonal and off-diagonal terms. This value depends on the strength of the draining effect and the number of cortical bins. Markuerkiaga et al. (2021b) derived a formula to adjust *p*2*t* = *p*2*t*_*n*_ to different bin sizes *n*. For this analysis, we used a value of *p*2*t*_3_ = 2.2, which corresponds to *p*2*t*_10_ = 6.3 determined there experimentally. To account for the uncertainty in the parameter, we show in the Supplementary Figure S3 that we obtain similar profiles for other realistic values of *p*2*t*. Using the deconvolved profiles in each subregion, we performed a repeated measures 2-way ANOVA, with factors *layer* (upper, middle, lower) and *attention condition* (attention to heartbeat, attention to sound). We tested for an interaction effect using an *F* -test.

## Acknowledgements

This study was supported by funding from the René and Susanne Braginsky Foundation (to KES), the ETH Foundation (KES), and the University of Zurich (KPP, KES).

## Appendix 1: Mask processing

### Transformation of masks

We transformed categorical masks with *N*_*cat*_ categories (for example the Brainnetome mask with 2*×* 6 insular subregions from MNI space to the individual subject space, defined by fMRI images) by the following algorithm:

1. For each of the *N*_*cat*_ categories, a binary 0-1 mask is transformed by the inverse warping field of the normalization of the MEWB image from MNI to the ME space.
2. The transformed (now continuous) masked are thresholded for each category at ≥ 0.5.
3. The resulting binary masks for each category are extended by 2 voxels (this is done in two steps: each time the mask is extended by the voxels which share a face with at least one voxel already contained in the mask).
4. A common mask over all categories is calculated by an voxel-wise logical *or* over the previous masks.
5. Within the common mask, each voxel is assigned to the category, whose mask from step 2 has the highest value (typically between 0 and 1, but could be higher/lower due to interpolation artifacts. We therefore took the absolute value before searching for the highest value to map slightly negative values to slightly positive values).
6. The procedure is repeated for the affine transformation between ME and fMRI space (without expansion of voxel).

### Correction of insular masks

We corrected the layer masks in section by the following algorithm to exclude potentially non-insular parts (like operculum). It is based on the heuristic, that in the insula, the lower layers are closer to the interhemispheric fissue than upper layers.

1. For each voxel *v* in the intersection of cortical ribbon mask and insular (Brainnetome) mask, let *u*_*v*_ denote the closest upper layer vertex (in the 75 % cortical-depth surface) and *l*_*v*_ the closest lower vertex (in the 25 % surface).^12^
2. Let 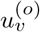 denote the opposite vertex in the *lower layer*, which corresponds to *u*_*v*_.^13^ Analogously, let 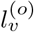 be the opposite vertex of *l*_*v*_ in the *upper layer*.^14^
3. If *v* is in the left hemisphere: We keep the voxel in the mask, if the *x*-coordinate of *l*_*v*_ is bigger than the *x*-coordinate of 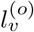 AND the *x*-coordinate of 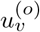 is bigger than the *x*-coordinate of *u*_*v*_. The same with *smaller than* if *v* is in the right hemisphere.
4. On the remaining mask, we keep the largest two^15^ connected components (determined by bwconncomp from MATLAB’s image processing toolbox with connectivity type 6 [voxels are connected if their faces touch]).

## Footnotes

1 Note that in contrast to the previous EEG-study Petzschner et al. (2019), where the sound was only played during the blocks of attention, in the current study the sound was played continuously during data acquisition. This was done to avoid potentially confounding effects due to interactions between auditory stimuli and the continuous presence of MR-scanner noise.

2 For this sequence, no field monitoring was performed. Images were directly reconstructed by the scanner.

3 But not resliced - so that only the header of *T*_1_ and fMRI images was modified.

4 The threshold was based on visual inspection of the blurring in the fMRI-images depending on the displacement w.r.t. the *B*_0_-map.

5 In the analysis plan of the preliminary analysis https://gitlab.ethz.ch/tnu/analysis-plans/muellerschraderetal_hbatt_ohbm we specified to smooth the contrast with a 3 mm FWHM Gaussian kernel. Since we decided to smooth the contrasts for the determination of functional masks with a 6 mm kernel (instead of 3 mm /compare the next footnote and the following discussion), we adapted this kernel accordingly.

6 In the analysis plan (Müller-Schrader et al., 2023), we specified to perform the functional masking based on first-level images, which were smoothed with a 3 mm FWHM Gaussian kernel. Later, we decided to use an (out-of-sample) second-level contrast to avoid statistical double dipping (compare footnote above). To account for between-subject variability, we increased the smoothing kernel to 6 mm.

7 Note that cortical depth is often looked at from the GM/CSF boundary downwards, so 0 cortical depth corresponds to that surface. Here, we used however surfaces, which were created by Freesurfer by first finding the GM/WM boundary and then inflating it. Therefore, 0 % corresponds here to the non-inflated GM/WM boundary, while 100 % corresponds to the fully-inflated GM/CSF boundary.

8 There was a correspondence between nodes in the lower and upper layers because Freesurfer determined the pial surface by inflating the GM/WM surface.

9 This procedure is symmetric, so it does not matter if we start by erasing *upper layer* or *lower layer* voxels.

10 We initially expected to observe an overlap of the functional masks with all 12 insular subregions and therefore specified a correction for 12 tests in the analysis plan. Since we now only performed three tests, we corrected only for these three.

11 Recall that they were computed by the equivolume principle.

12 We use the 25 % and 75 % layers, because they are smoother than the original (smoothed) surfaces, so we reduce the risk of accidentally removing a wrong voxel.

13 Recall that since Freesurfer inflates surfaces (and how the equivolume intermediate surfaces are computed), vertices at different cortical depths can be identified with each other.

14 Note that *u*_*v*_ and 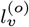 (which are both in the upper layer surface and close to *v* could be the same but don’t necessarily have to be.)

15 Note that we have both hemispheres in one single mask.

