## Supplemental Material for "Differences in layer-specific activation of the insula during interoceptive vs. exteroceptive attention"

#### 1 Results

##### 1.1 Non-layer analysis

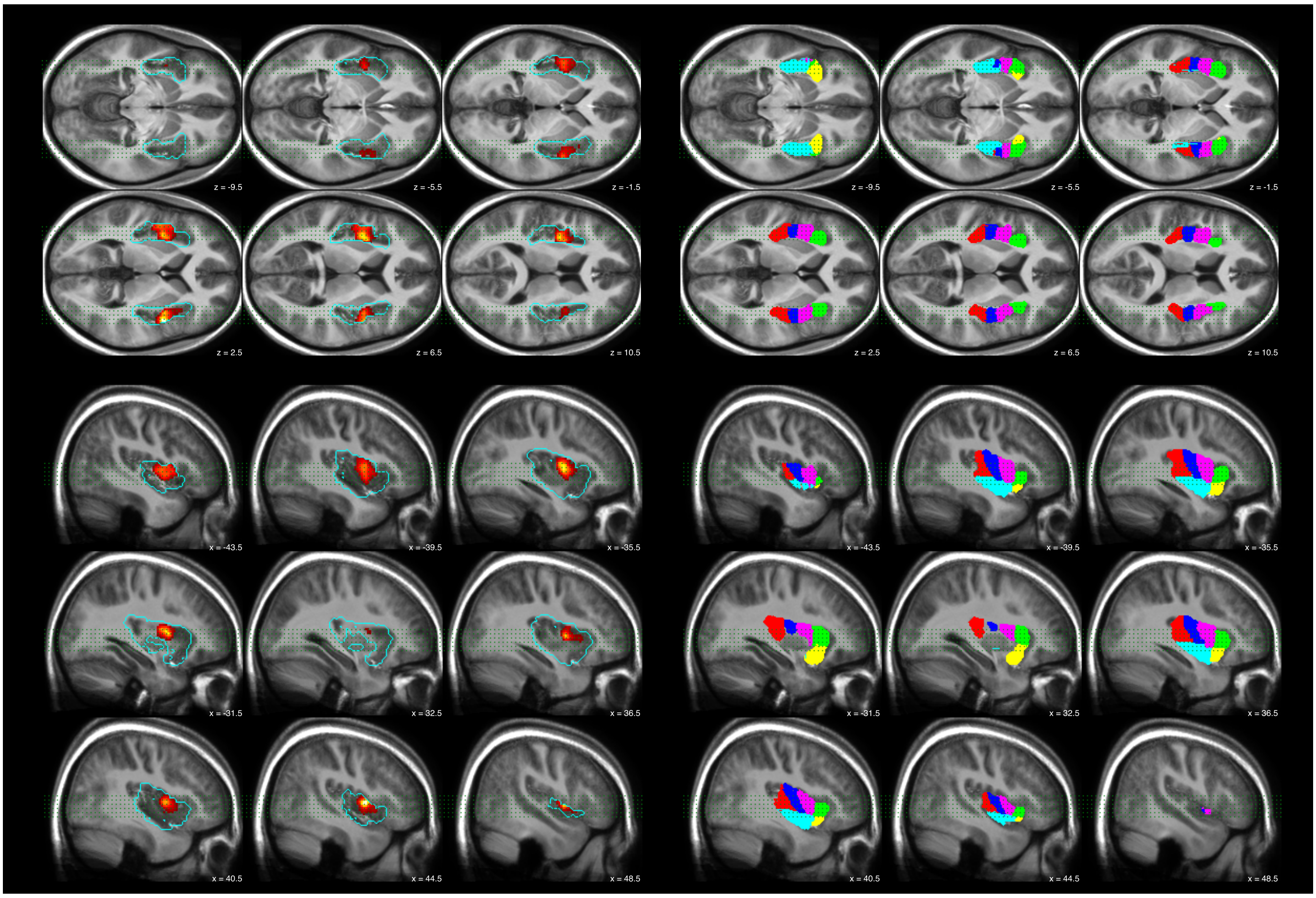


Figure S1: Results of non-layer analysis. Group level $F$-statistic of contrast $c_1 = \textit{attention to heart} - \textit{attention to sound}$ thresholded at cluster level (cluster defining threshold of $p = 0.001$). This is the same as what is shown in Figure 2, but for more slices. The thin green dots indicate the locations of the sagittal/transverse sections.

#### 1.2 ANOVA tables for layered analysis

##### 1.2.1 ANOVA tables using corrected layered mask (used for analyses)

| | Effect | DFn | DFd | F | p | p<.05 | ges |
|---|---|---|---|---|---|---|---|
| 1 | layers | 1.0 | 18.0 | 4.64 | 0.045 | * | 0.046 |
| 2 | conds | 1.0 | 18.0 | 1.25 | 0.278 | | 0.009 |
| 3 | layers:conds | 1.0 | 18.0 | 2.03 | 0.171 | | 0.014 |

Table S1: ANOVA table for layered analysis in left dorsal granular insula (**L__dIg**).

| | Effect | DFn | DFd | F | p | p<.05 | ges |
|---|---|---|---|---|---|---|---|
| 1 | layers | 1.0 | 18.0 | 0.58 | 0.456 | | 0.005 |
| 2 | conds | 1.0 | 18.0 | 42.95 | 0.000 | * | 0.238 |
| 3 | layers:conds | 1.0 | 18.0 | 15.63 | 0.001 | * | 0.023 |

Table S2: ANOVA table for layered analysis in left dorsal dysgranular insula (**L__dId**).

| | Effect | DFn | DFd | F | p | p<.05 | ges |
|---|---|---|---|---|---|---|---|
| 1 | layers | 1.0 | 18.0 | 7.83 | 0.012 | * | 0.088 |
| 2 | conds | 1.0 | 18.0 | 26.33 | 0.000 | * | 0.171 |
| 3 | layers:conds | 1.0 | 18.0 | 5.55 | 0.030 | * | 0.035 |

Table S3: ANOVA table for layered analysis in right dorsal dysgranular insula (**R__dId**).

##### 1.2.2 ANOVA tables using uncorrected layered mask (not used for analyses)

| | Effect | DFn | DFd | F | p | p<.05 | ges |
|---|---|---|---|---|---|---|---|
| 1 | layers | 1.0 | 18.0 | 1.70 | 0.209 | | 0.018 |
| 2 | conds | 1.0 | 18.0 | 7.15 | 0.016 | * | 0.066 |
| 3 | layers:conds | 1.0 | 18.0 | 1.10 | 0.309 | | 0.007 |

Table S4: ANOVA table for layered analysis using the **uncorrected** mask in left dorsal granular insula (**L__dIg**).

| | Effect | DFn | DFd | F | p | p<.05 | ges |
|---|---|---|---|---|---|---|---|
| 1 | layers | 1.0 | 18.0 | 1.31 | 0.268 | | 0.013 |
| 2 | conds | 1.0 | 18.0 | 42.26 | 0.000 | * | 0.258 |
| 3 | layers:conds | 1.0 | 18.0 | 25.60 | 0.000 | * | 0.047 |

Table S5: ANOVA table for layered analysis using the **uncorrected** mask in left dorsal dysgranular insula (**L__dId**).

| | Effect | DFn | DFd | F | p | p<.05 | ges |
|---|---|---|---|---|---|---|---|
| 1 | layers | 1.0 | 18.0 | 11.78 | 0.003 | * | 0.112 |
| 2 | conds | 1.0 | 18.0 | 37.72 | 0.000 | * | 0.233 |
| 3 | layers:conds | 1.0 | 18.0 | 12.61 | 0.002 | * | 0.047 |

Table S6: ANOVA table for layered analysis using the **uncorrected** mask in right dorsal dysgranular insula (**L__dIg**).

##### 1.2.3 Visualization of ANOVA for corrected/uncorrected mask

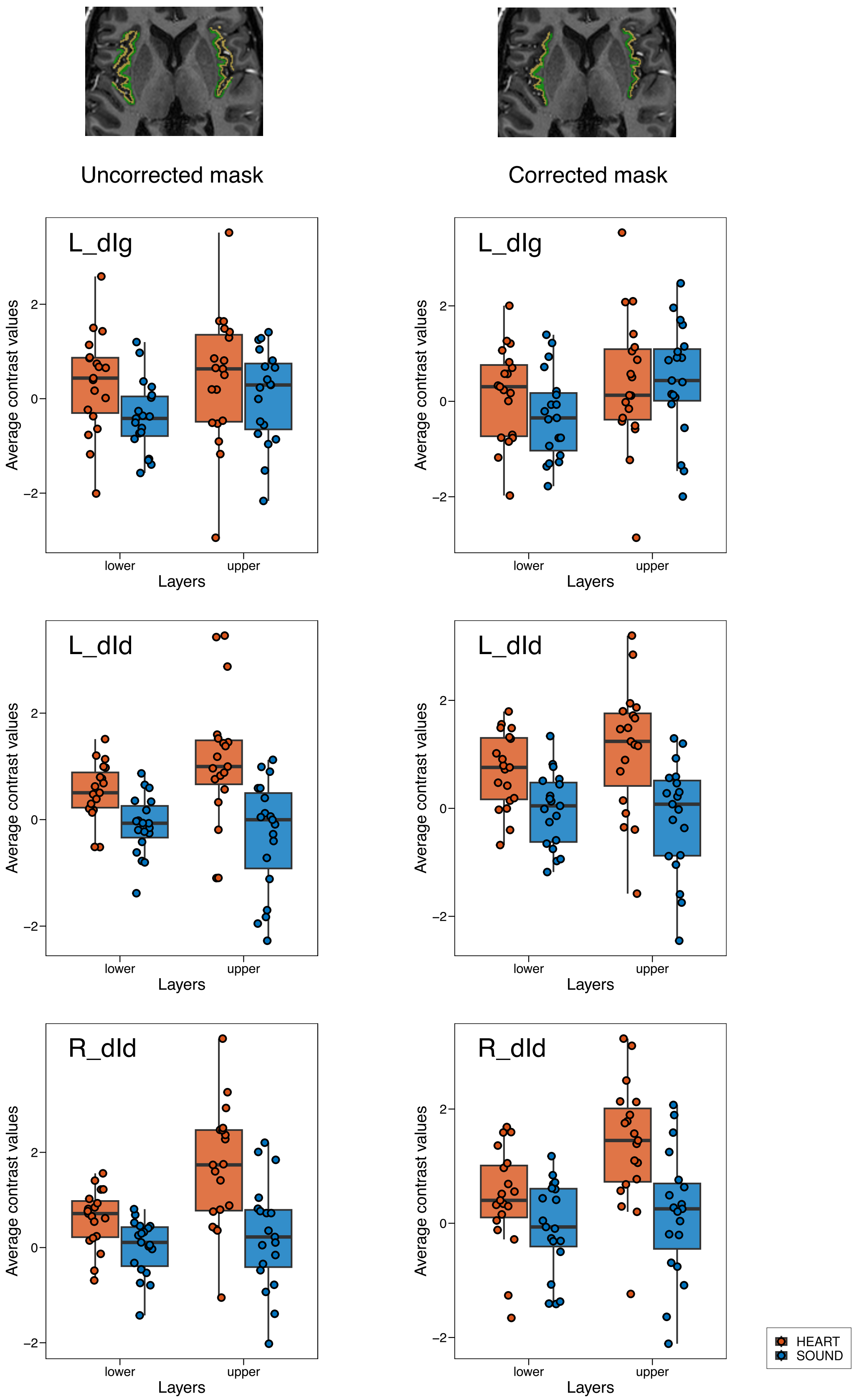


Figure S2: Visualization of within-subject ANOVA. The left column shows the results for the *original* masks, while the right column shows the results for the corrected masks (used in the analysis). The first row of plots shows the results for L_dIg, the second for L_dId and the third for R_dId.

#### 1.3 Layer profiles

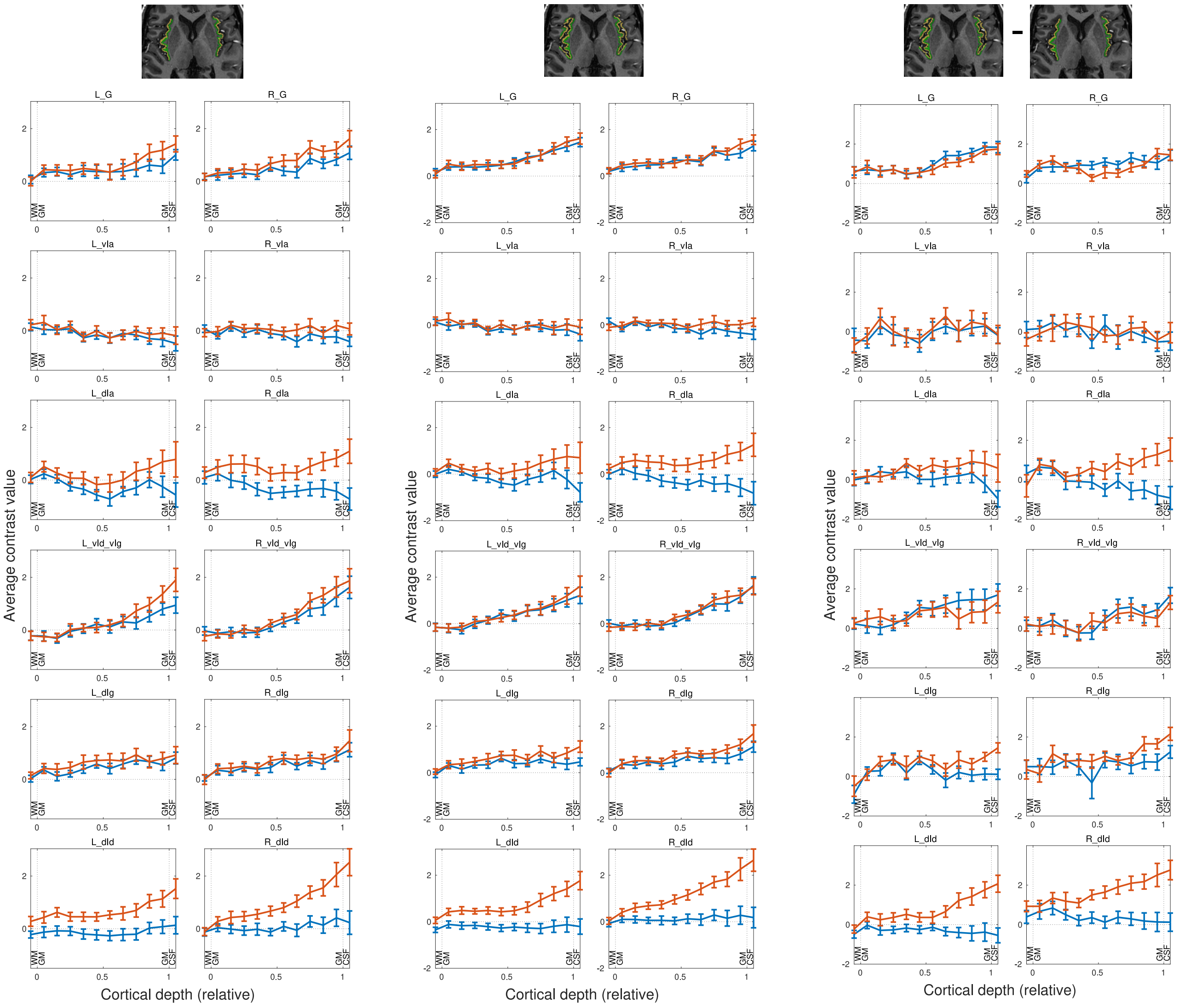


Figure S3: Layer profiles with the different masking strategies. Left: *Reduced* insular mask (used). Middle: *Original* insular mask. Right: Complement of *reduced* insular mask within *original* insular mask.

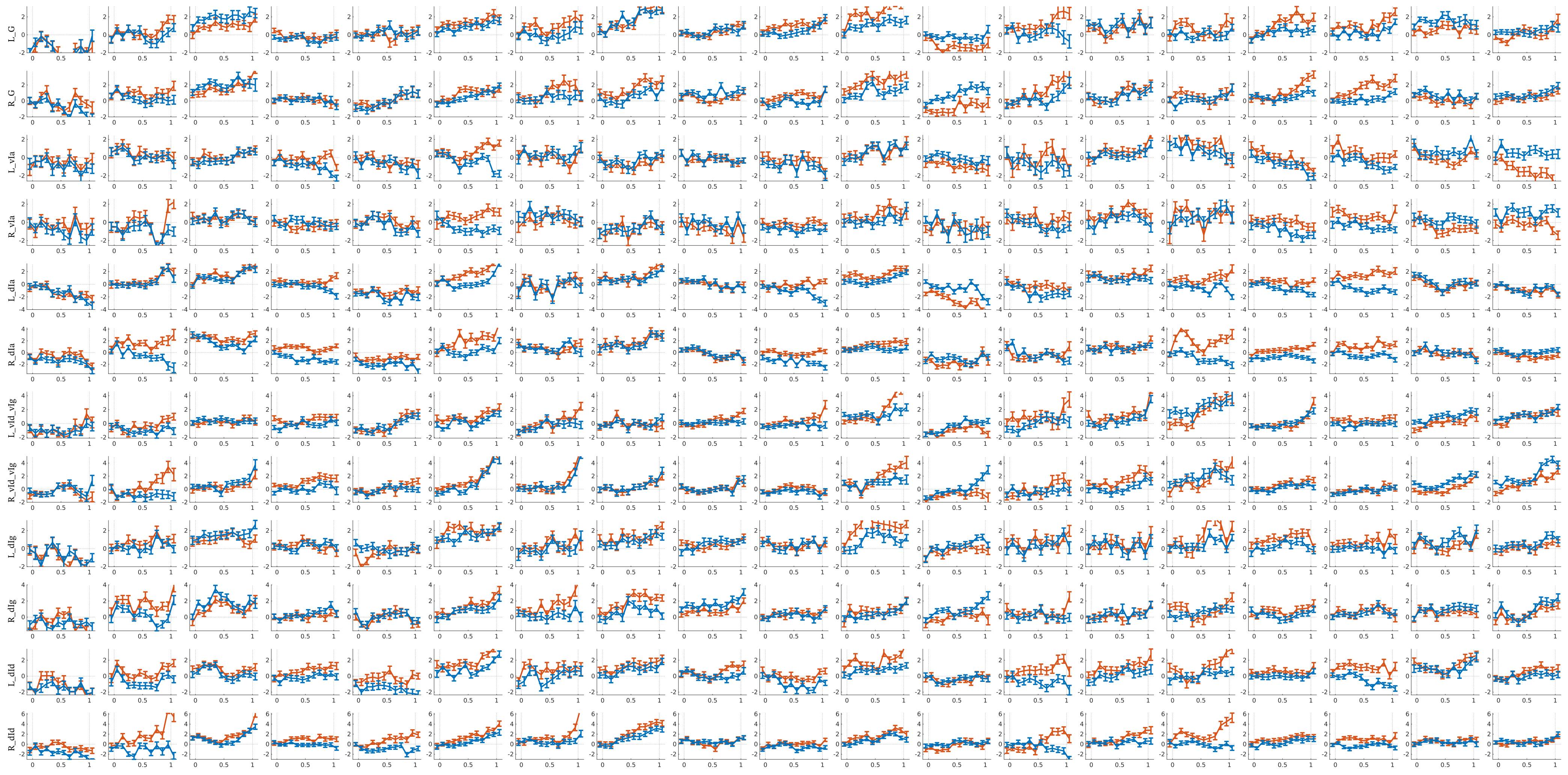


Figure S4: Single subject layer profiles. Rows: Insular subregions. Columns: Subjects.

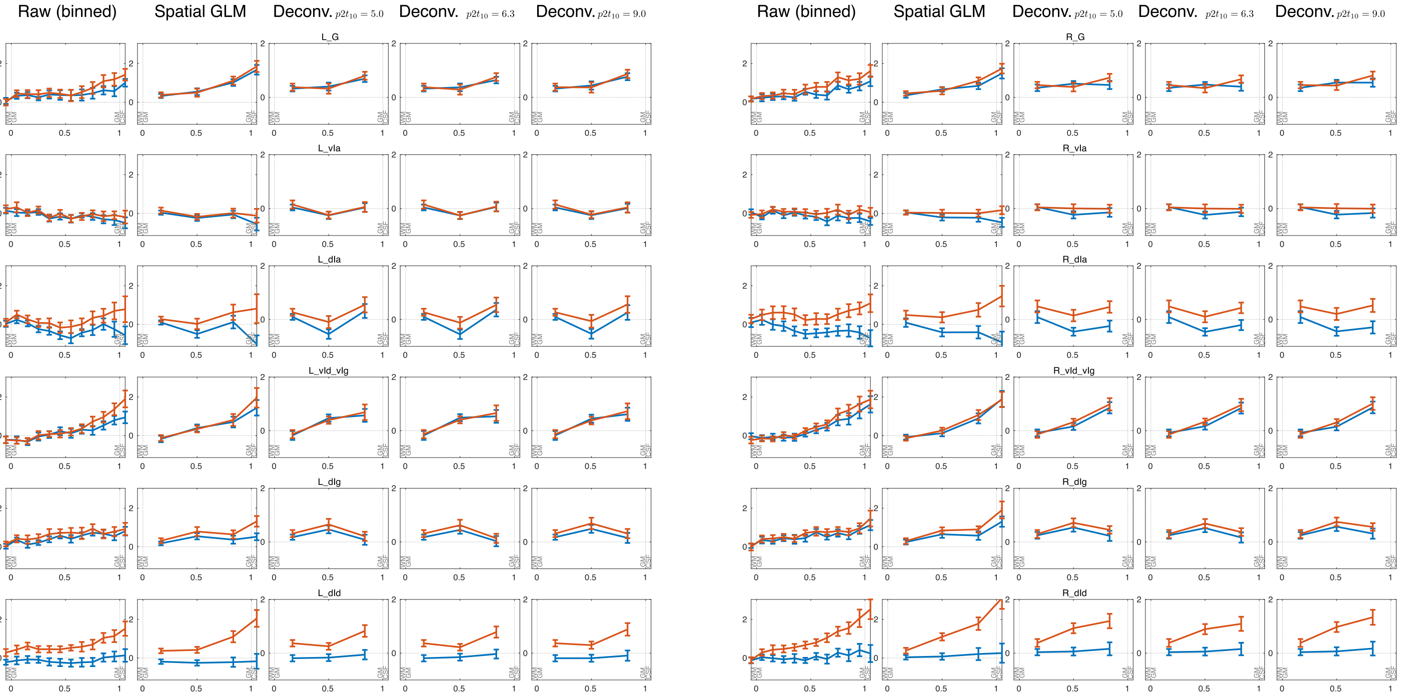


Figure S5: Results of (post-hoc) deconvolution of profiles for different values of $p2t$. The first three columns of each side are as in Figure 5, the fourth and fifth column show the results for deconvolution with different values of $p2t$.

### 2 Discussion

#### 2.1 Results from other (exteroceptive) layer attention studies

| Study | Attention cond. | Results | #Sub |
|---|---|---|---|
| [De Martino et al., 2015] | Sound in A1 | Task demands sharpen frequency tuning curves stronger in superficial cortical layers than in middle/deep layers | 5 |
| [Klein et al., 2018] | Attraction fields in V1 | Strongest population receptive fields (pRF) attraction in deep cortical portions (near the gray-white matter boundary) | 10 |
| [Gau et al., 2020] | Auditory and visual in auditory and visual cortices | Increased BOLD signal towards the cortical surface for auditory and visual conditions (but interpretation challenging) | 11 |
| [Liu et al., 2021] | Visual in V1 - V3 | Attention modulation stronger in superficial and deep (area V1) and superficial (areas V2,V3), respectively than middle layers | 15/12 |
| [Heynckes et al., 2023] | Auditory in primary and secondary auditory cortices | Detection of temporal shifts results in selective increase of activation in superficial layers | 10 |
| [van Mourik et al., 2023] | Visual in V1 - V3 | Increase in BOLD response, but no laminar-specific differentiation | 17 |

Table S7: Reported activation peaks from other layer fMRI studies on (exteroceptive) attention. The papers were obtained by searching the titles in the list of layer papers and preprints from `layermfri.com` for the word attention. Excluded: ”Dorsal Attention Network” and duplicates. The first column states the study, the second column the attention condition. The third column shortly summarizes the results, while the fourth column denotes the number of subjects included in the final analyses.

#### 2.2 Results from other interoceptive attention studies

Figure S6 and Table S8 compare the results obtained here with the results from other studies on interoceptive attention in the mid-insula.

| Study | Interoceptive attention | Distance | $x_{\mathrm{MNI}}$ | $y_{\mathrm{MNI}}$ | $z_{\mathrm{MNI}}$ |
|---|---|---|---|---|---|
| [Simmons et al., 2013] | Heart, stomach | 18 | 37 | -4 | 17 |
| | | 5 | 45 | 1 | 0 |
| [Avery et al., 2014] | Heart, stomach, bladder | 18 | 39 | -7 | 17 |
| | | 14 | -34 | -7 | 17 |
| [Avery et al., 2015] | Heart, stomach, bladder | 13 | -30 | 0 | 17 |
| | | 17 | 37 | 0 | 19 |
| | | 28 | 34 | -15 | 21 |
| | | 28 | 34 | -22 | 2 |
| | | 25 | -33 | -23 | 2 |
| [Farb et al., 2013] | Breathing | 31 | 33 | -18 | 21 |
| | | 22 | 33 | -6 | 18 |
| | | 23 | -33 | -15 | 21 |
| [Wang et al., 2019] | Breathing | 19 | -30 | 20 | 8 |
| | | 20 | 34 | 20 | 4 |
| This work | Heart | 0 | -36 | 2 | 6 |
| | | 0 | 46 | 4 | 4 |

Table S8: Reported activation peaks from other studies on interoceptive attention in the (mid) insula for interoceptive vs. exteroceptive attention. In case the peaks were clearly not in the mid insula (e.g. ventral anterior insula), they were not included in this list. Talairach coordinates were converted to MNI space using [Brett, 2004]. All coordinates and distances are reported in mm. Distances are with respect to the peak of activation determined in this work in the same hemisphere. The second column describes the interoceptive attention condition. See Figure S6 for a visualization.

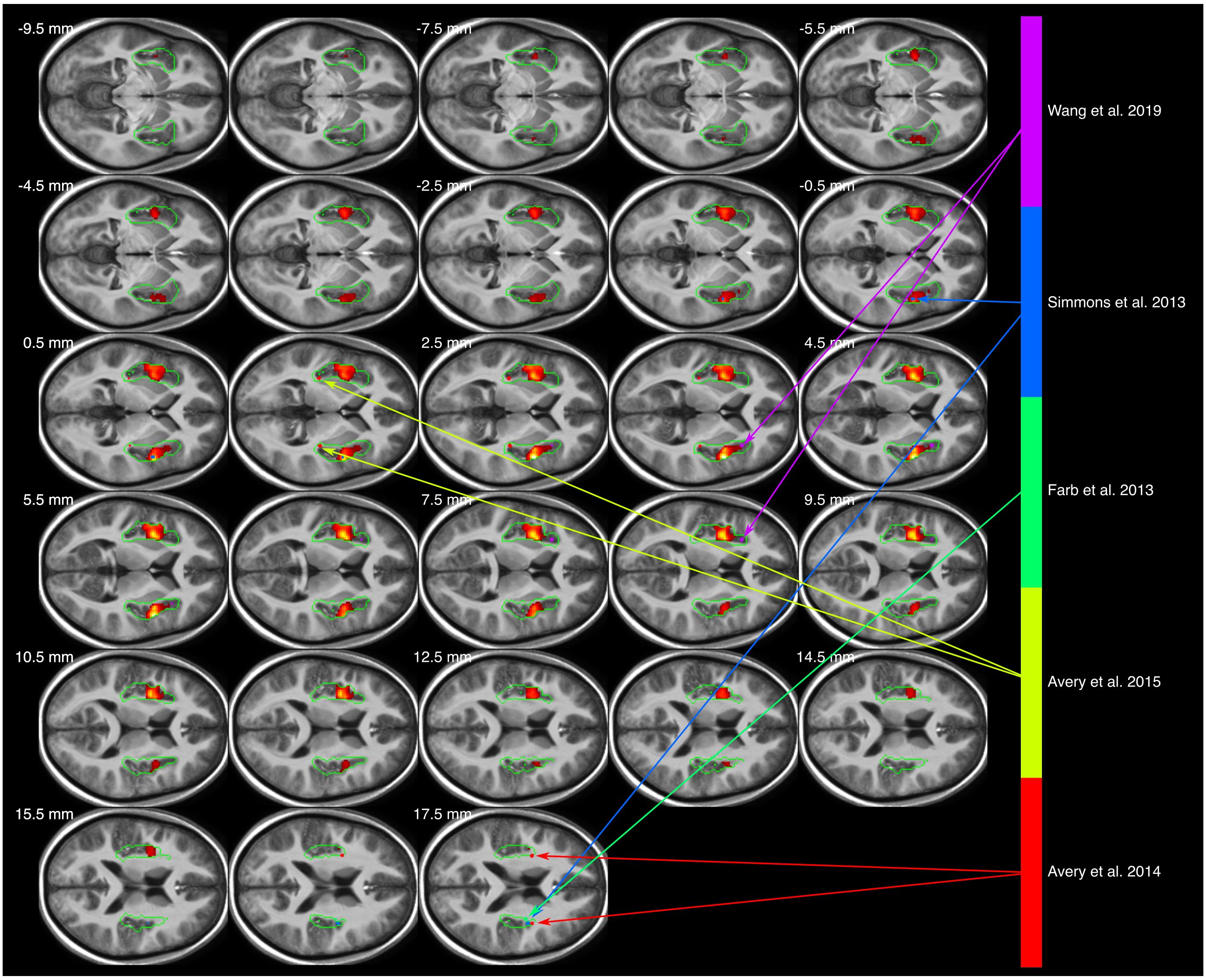


Figure S6: Visualization of $F$-statistic (thresholded at cluster level $p < 0.05$, cluster defining threshold $p < 0.001$; as in Figure 2) combined with peak coordinates of other studies. For list of peak coordinates see Table S8. The green mask is the intersection of insular mask and the field of view (FOV) for all subjects. The range of $z$-values is chosen from this range.
